# A bioelectric neural interface towards intuitive prosthetic control for amputees

**DOI:** 10.1101/2020.09.17.301663

**Authors:** Anh Tuan Nguyen, Jian Xu, Ming Jiang, Diu Khue Luu, Tong Wu, Wing-kin Tam, Wenfeng Zhao, Markus W. Drealan, Cynthia K. Overstreet, Qi Zhao, Jonathan Cheng, Edward W. Keefer, Zhi Yang

**Affiliations:** Biomedical Engineering, University of Minnesota, Minneapolis, MN, USA; Computer Science and Engineering, University of Minnesota, Minneapolis, MN, USA; Plastic Surgery, University of Texas Southwestern Medical Center, Dallas, TX, USA; Nerves Incorporated, Dallas, TX, USA; Fasikl Incorporated, Minneapolis, MN, USA

**Keywords:** artificial intelligence, deep learning, dexterous prosthetic hand, frequency-shaping amplifier, fully-integrated bioelectronics, human-machine interface, intrafascicular microelectrodes, intuitive control, neural decoder, peripheral neural pathways

## Abstract

**Objective:** While prosthetic hands with independently actuated digits have become commercially available, state-of-the-art human-machine interfaces (HMI) only permit control over a limited set of grasp patterns, which does not enable amputees to experience sufficient improvement in their daily activities to make an active prosthesis useful.

**Approach:** Here we present a technology platform combining fully-integrated bioelectronics, implantable intrafascicular microelectrodes and deep learning-based artificial intelligence (AI) to facilitate this missing bridge by tapping into the intricate motor control signals of peripheral nerves. The bioelectric neural interface includes an ultra-low-noise neural recording system to sense electroneurography (ENG) signals from microelectrode arrays implanted in the residual nerves, and AI models employing the recurrent neural network (RNN) architecture to decode the subject’s motor intention.

**Main results:** A pilot human study has been carried out on a transradial amputee. We demonstrate that the information channel established by the proposed neural interface is sufficient to provide high accuracy control of a prosthetic hand up to 15 degrees of freedom (DOF). The interface is intuitive as it directly maps complex prosthesis movements to the patient’s true intention.

**Significance:** Our study layouts the foundation towards not only a robust and dexterous control strategy for modern neuroprostheses at a near-natural level approaching that of the able hand, but also an intuitive conduit for connecting human minds and machines through the peripheral neural pathways.

**Clinical trial:** DExterous Hand Control Through Fascicular Targeting (DEFT). Identifier: NCT02994160.

## 1. Introduction

Neuroprosthetics is a scientific discipline that has been interestingly advertised in science fiction movies. RoboCop, Terminator, and Avatar are examples that present vivid views on what would happen if the human minds and machines can efficiently communicate: a human-machine symbiosis. DEKA Corp. has embodied the connection between the Star Wars universe and real-world devices with the production of the Luke arm [41]. Though exaggerated, a combination of mind-reading capability and robotics can benefit many people and society at large. For example, neuroprosthetics research can benefit millions of people with motor impairment, including the amputee population as well as those who have neural injuries in the spinal cord and the brain. Among different applications, the use of neuroprosthetics to restore lost function in upper limb amputees is the most challenging scenario. From a mechanistic perspective, the human hand is the most complex device we directly control, and the hand is innervated by the largest number of sensory neurons of any organ in the body.

The development of upper limb prostheses has been benefitted from progress in robotics and material science, where the newer designs such as the DEKA Arm [42, 43], the APL ARM [29], and the DLR Hand Arm system [16, 17] can support a wide range of movements. However, none of them can be fully utilized due to “ineffective control”. What is missing is an human-machine interface (HMI) that can decode the motor intention in the brain, and enable intuitive, simultaneous control of all available degrees-of-freedom (DOF) of the prostheses with lifelike dexterity [6, 48].

In the literature, there are three locations where motor control signals can be intercepted: the brain [2, 22, 26, 70], muscles [10, 11, 18, 28, 33], and peripheral nerves [1, 8, 27, 46, 53]. While cortical decoding techniques with implanted microelectrode arrays in the brain have pioneered the research field for many years, it remains unclear if there could be sufficient neural information harvested to meaningfully restore the lost motor function. For example, it is not yet possible to demonstrate near-natural, individual finger control with a brain implant.

Prosthetics based on surface electromyography (EMG) signals recorded from available muscles in the amputated limb are non-invasive and have been widely adopted by amputee patients [10, 45]. However, these EMG prosthetics, regardless of the allowable DOF in the arm system, only permit sequential control of grasp patterns such as opening and closing the prosthesis through the co-contraction of residual muscle sites. Higher DOF control is possible with pattern recognition systems that may provide 3-4 simultaneous DOF control, but remain non-intuitive, unnatural, and cannot be generalized in daily tasks [19]. With targeted muscle reinnervation (TMR) intuitive control of some prosthetic functions may be possible, but requires a significant surgery, and is intrinsically unpredictable in outcome [3].

This work focuses on motor decoding using electroneurography (ENG) signals acquired from peripheral nerves. There are multiple challenges associated with this approach [5, 24, 36, 47, 54]. For example, typical ENG signals acquired from extra-and-intraneural electrodes have an amplitude ranging from a few to tens of microvolts, which are orders of magnitude smaller than EMG and cortical neural signals. ENG recordings are also contaminated by large-amplitude interferences, such as those originating from body motions and residual muscles. Furthermore, ENG signals obtained from one electrode are often combined activities from hundreds of individual axons. As a result, a substantial level of processing and pattern recognition is required to isolate and extract the desired data features. More recent studies on microelectrodes [4, 40], bioelectronics [67], AI algorithms [9], and clinical validations [7, 12, 39, 55, 69] argue multi-DOF motor control via the peripheral neural pathways may be a more effective and promising approach.

Fig. 1 illustrates an artist’s concept of the proposed neural interface, which generally consists of three components: the intrafascicular microelectrodes, the fully-integrated bioelectronics, and the artificial intelligence (AI) decoder. The longitudinal intrafascicular electrodes (LIFE) microelectrode arrays are implanted into the median and ulnar nerves of the amputee using microsurgical fascicular targeting (FAST) technique, creating the interface with individual nerve fascicles. Nerve signals or ENG are acquired by ultra-low-noise neural recording chips based on the frequency-shaping technique we have pioneered [67]. The result is a continuous stream of neural data, in which the patient’s movement intentions are encoded. Next, we design an AI model based on recurrent neural network (RNN) architecture to perform regressive motor decoding of 15 DOF that includes the flexion/extension and abduction/adduction motions of all five fingers, thus proving a definite proof that the contents of this information channel are sufficient for intuitive, dexterous control of a prosthetic hand.

**Figure 1.**
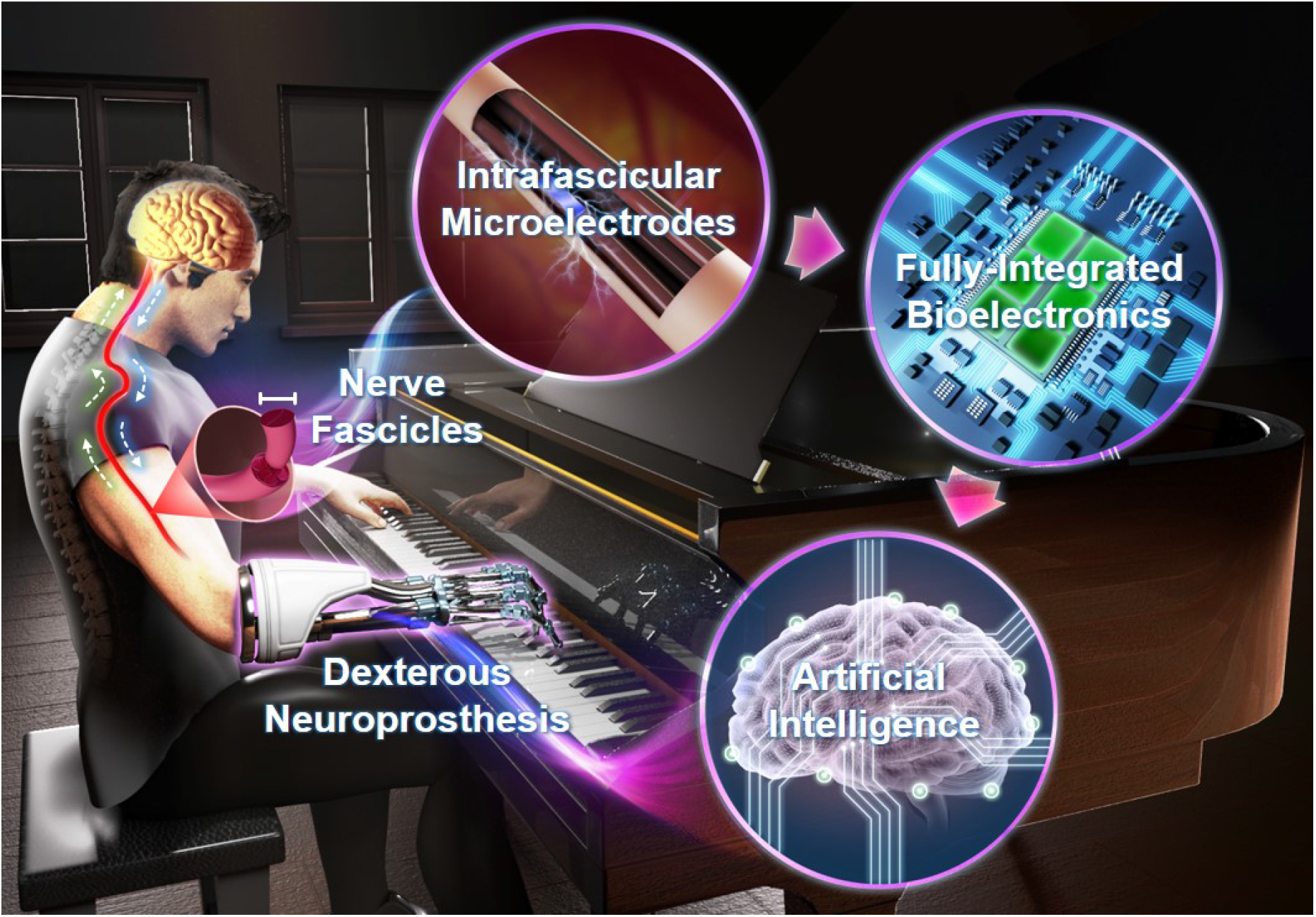
An artist’s concept of the proposed neural interface for dexterous neuroprostheses. The nerve technology platform presented here combines fully-integrated bioelectronics, intrafascicular implantable microelectrodes and deep learning-based artificial intelligence (AI) to facilitate an intuitive connection between human intent and dexterous neuroprostheses.

The organization of the paper is as follows. Section 2 describes the implementation of the proposed neural interface including the Scorpius neural recording system, LIFE-FAST microelectrodes, and design of the AI model. Section 3 presents the measurements of ENG signals and benchmarking of the AI model. Section 4 provides discussions about the results and future directions. Section 5 concludes the paper.

## 2. Methods

### 2.1. Experimental Paradigm & Protocol

An overview of the experimental setup is presented in Fig. 2(A,B). A transradial amputee subject is implanted with four microelectrode arrays targeted to four different fascicles within the median and ulnar nerves. The mirrored bilateral paradigm is utilized where ENG signals are recorded from the injured hand, while the intended movements are captured by the data glove from the able hand (ground truth). The acquired ENG dataset with ground-truth labels is used to train AI models that can report the subjects’ motor intention.

**Figure 2.**
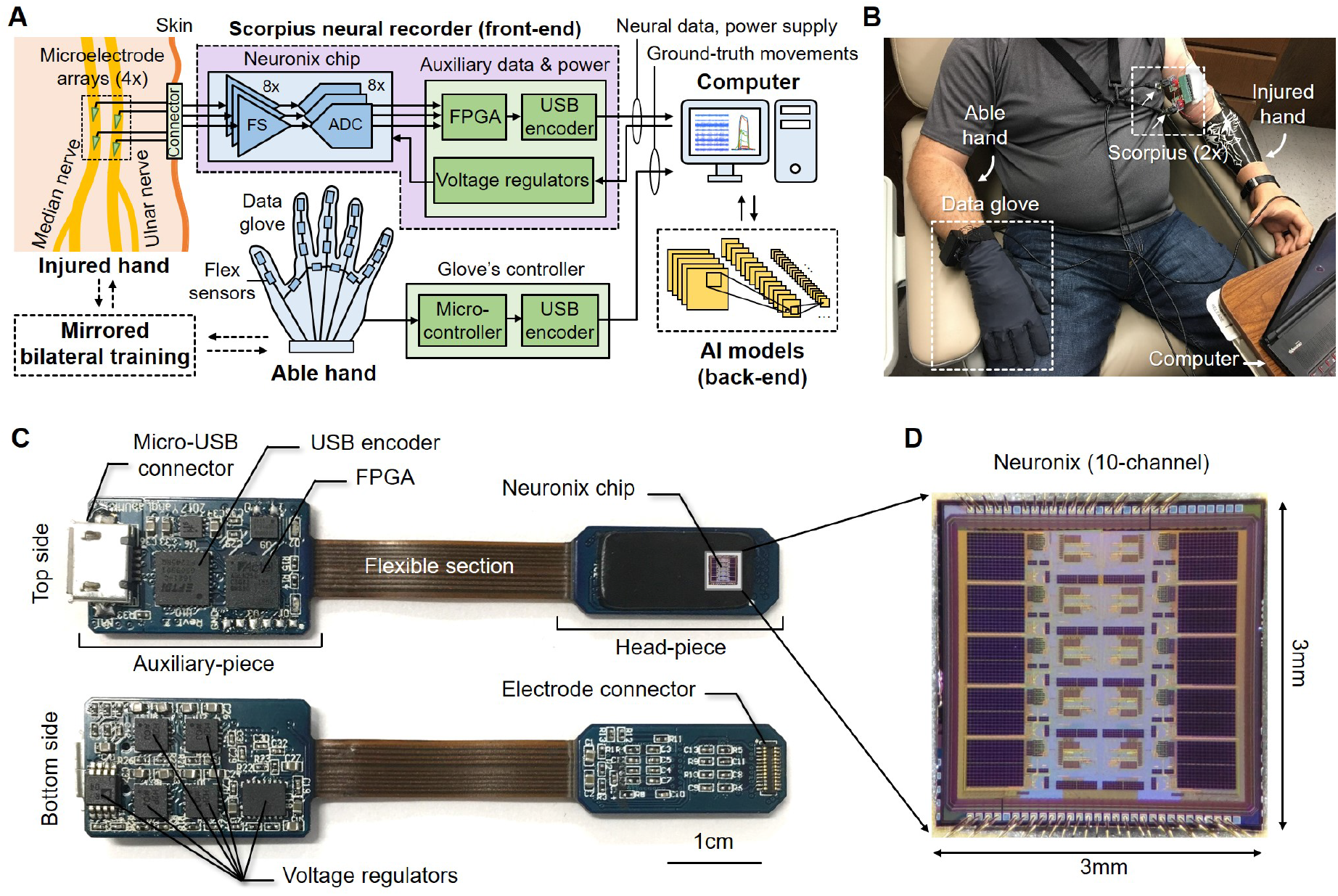
Experimental paradigm and hardware components. (A) Diagram of the experiment setup including the Scorpius front-end neural recording system, implanted FAST-LIFE microelectrode arrays, a computer for data acquisition and back-end AI models implementation, and a data glove for obtaining ground-truth movements during mirrored bilateral training. (B) Photo of the transradial amputee during an experiment session (C) The Scorpius is a miniaturized prototype designed toward implantable applications. The head-piece contains a fully-integrated Neuronix neural recording chip, an electrode connector, and passive components while the auxiliary-piece includes voltage regulators and data relaying circuits. (D) Microphotograph of the Neuronix chip that has ten fully-integrated, high-performance neural recording channels based on the frequency-shaping (FS) architecture.

The human experiment protocols are reviewed and approved by the Institutional Review Board (IRB) at the University of Minnesota (UMN) and the University of Texas Southwestern Medical Center (UTSW). The patient voluntarily participates in our study and is informed of the methods, aims, benefits and potential risks of the experiments prior to signing the Informed Consent. Patient safety and data privacy are overseen by the Data and Safety Monitoring Committee (DSMC) at UTSW. The implantation, initial testing, and post-operative care are performed at UTSW by Dr. Cheng and Dr. Keefer while motor decoding experiments are performed at UMN by Dr. Yang’s lab. The clinical team travels with the patient in each experiment session.

### 2.2. The Scorpius Neural Recording System

#### Scorpius

Fig. 2(C) shows the overview of the Scorpius system – a miniaturized front-end recorder equipped with the fully-integrated Neuronix neural recording chip. It consists of two sub-units: the head-piece and the auxiliary-piece, connected by a flexible section. The head-piece contains the Neuronix chip - the recorder’s key functional component, along with an electrode connector, and other passive components. The auxiliary-piece consists of a field-programmable gate array (FPGA) (AGLN250, Microsemi, CA), a high-speed universal serial bus (USB) encoder (FT601Q, FTDI, UK), and power management circuitry with various voltage regulators (ADR440, ADP222, ADA4898-2, Analog Devices, MA). The auxiliary-piece sole function is to pass-through the digitized neural data while powering the Neuronix chip through a single micro-USB connector. Recorded data are retrieved and processed offline on the back-end on an external computer.

#### Neuronix

members of the Neuronix chip family are a fully-integrated neural recorder application-specific integrated circuits (ASIC) based on the frequency-shaping (FS) architecture [58, 59, 61–63, 65–67] and high-resolution analog-to-digital converters (ADC) [38, 56, 60, 64] that we have pioneered. At the sampling rate of 40 kHz, the recorder is optimized to acquire neural data with a maximum gain of 60 dB in the bandwidth 300-3000 Hz and approximately 40 dB in the bandwidth 10-300 Hz while suppressing motion and stimulation artifacts. This frequency-dependent transfer function is achieved with the switched-capacitor circuit implementation, which offers a minimal noise floor while simultaneously preventing input circuit saturation due to large-amplitude artifacts. The FS architecture’s unique characteristics allow the Neuronix chip to capture high-quality neural signals by “compressing” the effective dynamic range (DR) of the recorded data, giving an additional 4.5-bit resolution across the input full-range as shown in Xu *et al*. [61]. This is coupled with a 12-bit successive approximation register (SAR) ADC with automatic calibrations including comparator’s random offset cancellation and capacitor mismatch error suppression. The resulted system has a “11+4.5” bits effective DR, where 11-bit is the effective number of bits (ENOB) of the ADC and 4.5-bit is the DR compression ratio. These capabilities have been extensively validated and demonstrated in various Neuronix chip generations, in both in-vitro and in-vivo experiments [58, 59, 67]. Here we solely focus on the system integration where Neuronix is deployed for human clinical applications. The latest iteration of the Neuronix chip shown in Fig. 2(D) has ten recording channels with a further optimized layout that occupies a 3mm x 3mm silicon die area when fabricated in the GlobalFoundries 0.13*μ*m standard CMOS process. Eight channels are used for neural data acquisition and two channels are reserved for testing purposes.

Table 1 summarizes the specifications of the Scorpius system in comparison to commercial counterparts [34, 44, 51]. Here we only include systems that are being used in human clinical trials for neuroprosthesis studies involving peripheral nerves or cortical recordings. The Scorpius has a substantially smaller footprint (size and weight) compared to others because it is the only miniaturized system specifically designed towards implantable and wearable applications while showing improved sensitivity and specificity in sensing nerve signals. Furthermore, the FS architecture allows the proposed system to achieve an effective system resolution of “11+4.5” bits which is significantly higher than the commercial counterparts. In practice, the overall system resolution is the effective DR which is determined by the input range, noise floor, and ADC resolution. A combination of a higher system resolution as well as system miniaturization and low power implementation allows Scorpius to isolate weak neural signals from large amplitude artifacts and interferences, which in turn enable the high DOF motor decoding through deep neural networks.

**Table 1.** Specifications of Scorpius in comparison to commercial systems being used in human clinical applications.

|  | Cerebus/NeuroPort<br>(Blackrock Microsys.) | Grapevine<br>(Ripple Neuro) | RHD2000<br>(Intan Tech.) | Scorpius<br>(This work) |
| --- | --- | --- | --- | --- |
| Die size (mm) | - | - | 4.1 x 4.8 | 3 x 3 |
| System size<sup>1</sup> (mm) | 88 x 325 x 425 | 43 x 110 x 190 | 20 x 99 x 155 | 4 x 12 x 68 |
| System weight (g) | 6800 | 907 | 130 | 3 |
| No. of channels<sup>2</sup> | 8-512 | 32-512 | 16-256 | 10 |
| Sampling rate (kHz) | 30 | 30 | 30 | 10-80 |
| Bandwidth (Hz) | 0.3-7,500 | 0.1-7,500 | 0.1-10,000 | 10-5,000 |
| Noise floor (μV<sub>rms</sub>) | 3.0 | 2.1 | 2.4 | 2.5 |
| ADC resolution (bits) | 16 | 16 | 16 | 12 |
| System resolution<sup>3</sup> (bits) | 12.4 | 12.5 | 12.0 | 11+4.5 |
| Neuroprosthesis<br>human clinical trials | Davis et al. (2016)<br>Irwin et al. (2017)<br>Vu et al. (2018) | Wendelken et al (2017)<br>George et al. (2018) | Bullard et al. (2019)<br>Kannape et al. (2016) | This work |

<sup>1</sup>System size & weight includes the headstages and power/data interface board.
<sup>2</sup>No. of channels depend on the number of headstages connected (1-8).
<sup>3</sup>System resolution is the effective dynamic range (DR) that is determined by the input range, noise floor, and ADC resolution.

### 2.3. FAST-LIFE Microelectrode Array

Fig. 3 shows the construction of the microelectrode array that is an extended version of the design reported in [4, 40]. Each array consists of four cuff contacts and an intrafascicular shank with ten longitudinal intrafascicular electrodes (LIFE) contacts spaced 0.5mm apart. The design aims at recording neural data from both single-axon and population of neurons as well as providing sensory feedback through electrical stimulation.

**Figure 3.**
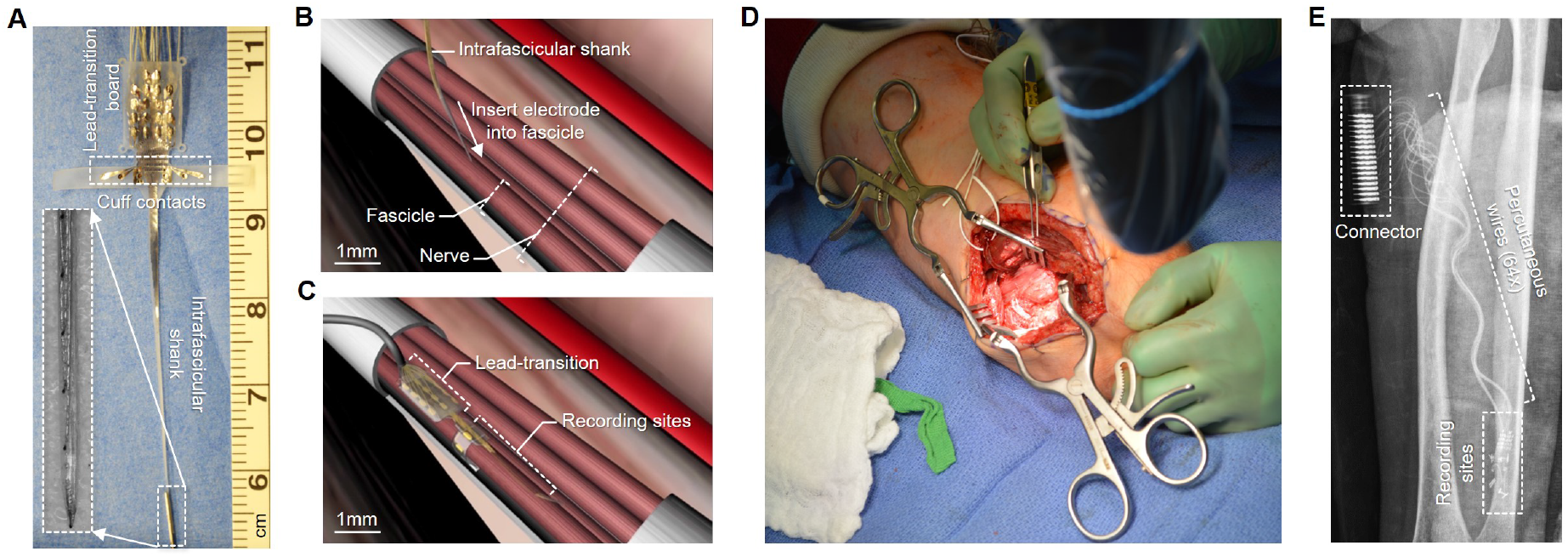
FAST-LIFE microelectrode array design and implantation site. (A) Photo of the proposed microelectrode array which consists of cuff electrodes and an intrafascicular shank with ten electrode contacts. (B) Illustration of the implantation procedures. (C) Illustration of the implanted electrodes and recording sites. (D) Photo of the implantation sites during surgery. The human subject is implanted with four arrays in four different fascicles, two in the median nerve and the other two in the ulnar nerve. (E) X-ray image of the patient’s arm after the operation.

The implantation surgery is performed by Dr. Cheng at the Clements University Hospital in UTSW, Dallas. Microsurgical fascicular targeting (FAST) technique is used to guide the microelectrode array into the nerve fascicle as shown in Fig. 3(B, C). The patient is implanted with four FAST-LIFE microelectrode arrays, two in the median nerve, and the other two in the ulnar nerve. Within one nerve, the arrays are placed into two discrete fascicle bundles. Their functional specificities are examined during the surgery such that one is more likely to carry efferent motor control signals while the other carries the afferent sensory/proprioceptive signals. The “motor” fascicles are identified by using a handheld nerve stimulator (Vari-Stim III, Medtronic, MN) to induce contraction of intrinsic muscles still present in the residual limb. The “sensory” fascicles are found by placing needle electrodes in the scalp over the primary sensory cortex and observing somatosensory evoked potentials (SSEP) elicited by the nerve stimulation. Nevertheless, it is worth noting that there is no clear separation between motor and sensory functions as shown in [40]. An electrode with strong motor control signals can frequently be found on the “sensory” fascicles and vice versa.

Fig. 3(D) shows a photo of the implantation site during the operation. Fig. 3(E) shows an x-ray image of the patient arm after the operation with the implanted microelectrode arrays, percutaneous wires, and the external connector. In total, 56 wires are brought out through 14 percutaneous holes where each hole supports a 4-wire bundle. One of the cuffs is left out and replaced with a platinum wire that acts as the common ground and return-electrode for electrical stimulation. The external wires are attached to two connector blocks which can be connected to external recorders via two standard 40-pin Omnetics nano connectors.

### 2.4. Mirrored Bilateral Training

Motor decoding data are obtained via the mirrored bilateral training with a data glove, which has been successfully implemented in similar studies [28, 49]. The patient is asked to perform various gestures with the able hand while imagining doing the same movement simultaneously with the injured (phantom) hand. Nerve data from the injured hand are acquired from implanted microelectrodes while the intended motions from the able hand are concurrently captured by a data glove (VMG 30, Virtual Motion Labs, TX). The glove can acquire 15 DOF corresponding to the movement of key joints as shown in Fig. 4(A). They include the flexion/extension of all five fingers (D1-D10), abduction/adduction (D11-D14), and thumb-palm crossing (D15). The glove’s data are used as the ground-truth label for both training and benchmarking the deep learning model.

**Figure 4.**
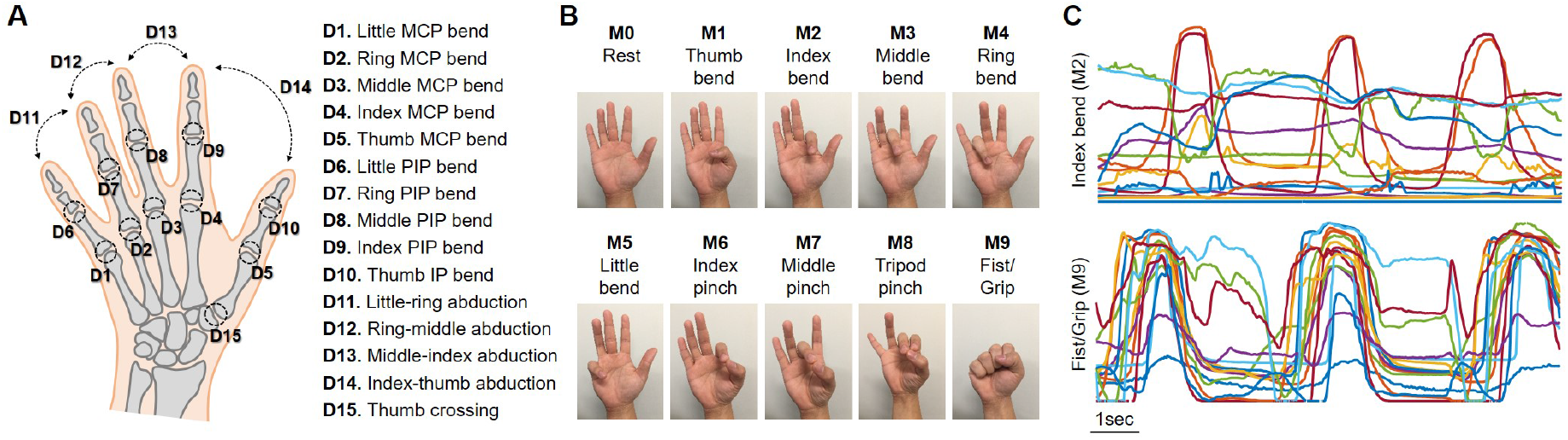
Mirrored bilateral paradigm is used to collect nerve data with ground-truth labels of intended movements. (A) Map of 15 DOF captured by the data glove and their corresponding joints on the hand. (B) Illustration of different hand gestures used for training. The gestures are selected to elicit at least two or more DOF simultaneously. (C) Example of intended movement trajectory of two different gestures: M2 (index bend) where only 4-5 DOF corresponding to one finger are strongly elicited, and M9 (fist/grip) where almost all DOF are excited to some degree.

While it may be appealing to be able to demonstrate independent and exclusive control of 15 individual DOF, there are several practical challenges to this approach.

- (i) It technically requires training and testing all 2^15^ combinations of all the DOF, which is not possible.
- (ii) In daily living activities, most people only use a small fraction of these possible gestures which almost always involve the movement two or more DOF.
- (iii) Depending on the person, some DOF cannot be even independently controlled; for example, many people struggle to flex the little finger without also bending the ring finger.

In fact, the coherent motion of multiple DOF contributes to the intuitiveness and dexterity of the human hand that cannot be easily replicated by a mechanical prosthesis. Here we must accept the trade-off between practicality and generativity by starting with the most common hand gestures such as flexing fingers, grasping, pinching, etc. In future studies, additional gestures could be added like thumb up, pointing, etc. Thanks to the robustness of deep learning, this can easily be done by adding more data into the existing dataset and fine-tuning the models without any other modifications to their architecture.

Fig. 4(B) illustrates different hand gestures utilized in this experiment which are selected to elicit at least two or more DOF at once to various degrees. They comprise of individual finger flexion (M1-M5), two or three fingers pinching (M6-M8), and fist/gripping (M9). Resting (M0) is used as the reference position. In each experiment session, the patient repeatedly alters between resting (M0) and one of the other hand gestures (M1-M9) at least 100 times where neural data and intended motions are continuously recorded. Incorrect trials are manually discarded upon visual inspection. They mostly consist of trials where the burst of neural activities do not synchronize with the movement ground-truth due to the inconsistent timing of the data glove. The final dataset which is used to train the AI models has a total of 760 trials. Fig. 4(C) shows typical examples of the raw data acquired by the data glove during the experiment. In certain gestures such as index bend (M2), only 4-5 DOFs out of 15 DOF are strongly excited, while in others such as fist/grip (M9), almost all DOF are elicited to some degree.

### 2.5. Deep Learning-Based AI Models

Fig. 5(A) presents an overview of the back-end data analysis paradigm including pre-processing procedures and motor decoding using deep learning. In this work, the data are processed offline with the primary goals of (i) demonstrating ENG recordings obtained from the proposed Scorpius system contain sufficient information for multi-DOF motor decoding; and (ii) determining the best machine learning approach to extract and interpret this information.

**Figure 5.**
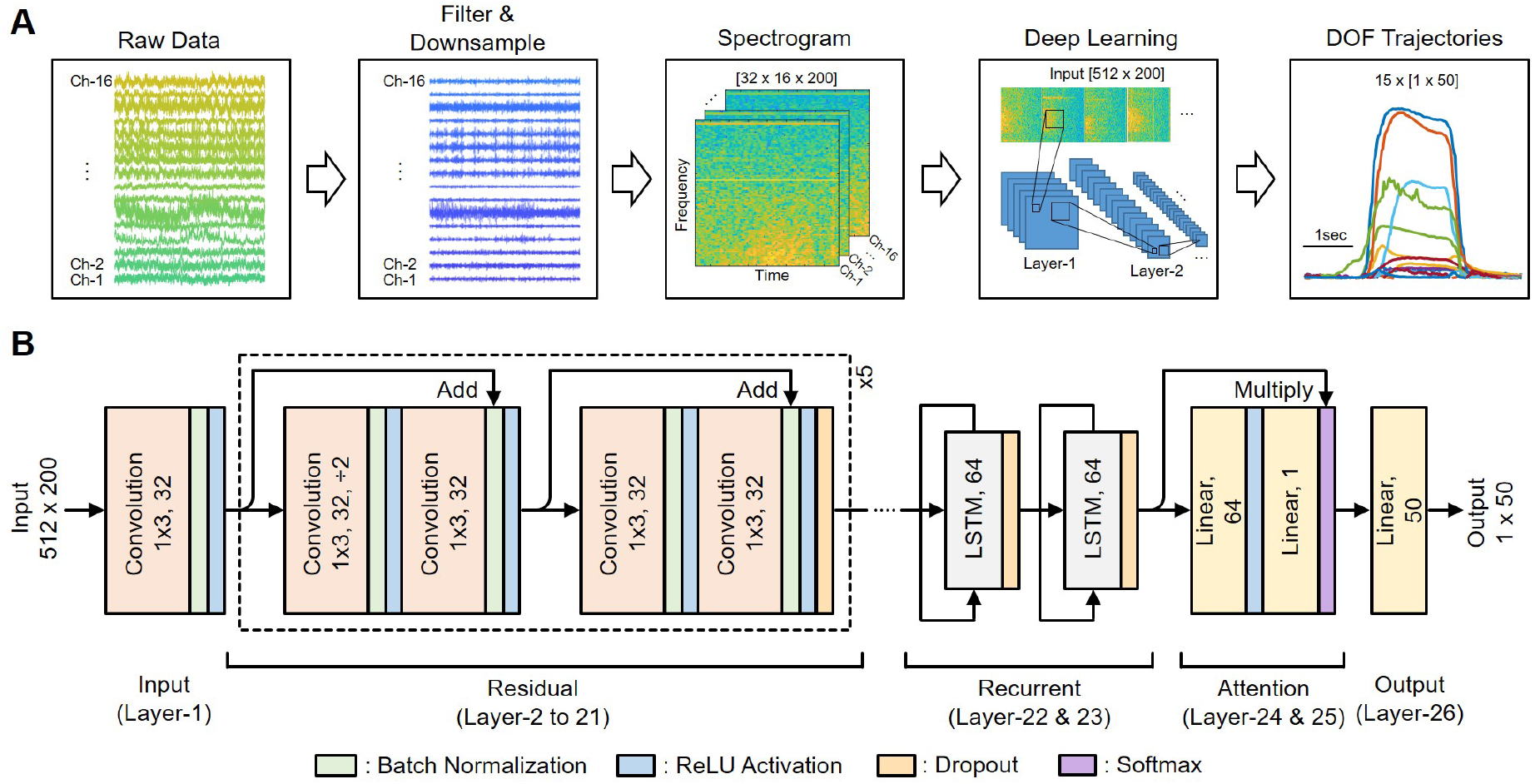
Overview of the back-end signal processing and deep learning-based AI decoders. (A) Diagrams of the signal pre-processing flow including filtering, features extraction via spectrogram, and motor decoding using deep learning-based AI models. (B) Design of the AI model which is based on a 26-layer recurrent neural network (RNN) architecture including residual, recurrent and attention blocks. A total of 15 AI models with the same structure are trained for every DOF. Each model takes the input from all 16 recording channels.

#### Pre-processing

The recorded data including neural signals from the Scorpius system and the labeled ground-truth movements from the data glove are segmented into individual, non-overlapping trials of approximately four seconds. The raw neural data are first pre-processed by applying anti-aliasing filters and downsampling four times. Next, neural features are extracted using the short-term Fourier transform (spectrogram) with 20 ms steps. For each channel, we compute the average power spectral density of 32 frequency bins spaced on a log scale from 150 to 3000 Hz. We choose a conservative cut-off frequency of 150 Hz to capture most of the ENG signals’ power while minimizing the EMG signals and motion artifacts that may occur during the experiment and tend to locate in the lower frequency band. This results in an input of dimension [16 channels] x [32 bins] x [200 time-steps], which is reshaped into an [512 x 200] “image” for the deep learning model. Here spectrogram serves as feature extraction and dimensionality reduction which significantly lower the complexity of the AI decoder. While it is possible to design deep neural networks that directly use raw neural data, the number of parameters/layers and computational resources needed would be considerably higher making it further difficult to translate into real-world applications.

#### Deep learning model

Fig. 5(B) presents the design of the core AI decoder which is based on the RNN architecture with the use of long-short term memory (LSTM) cells. The RNN architecture is among the most advantageous approaches for handling sequential modeling problems [15, 23, 52, 68]. Here we developed a self-attention mechanism with the RNN to encode the recorded high-dimensional signals into a 64-dimensional neural code that can be used to generate the motion trajectories. The network architecture comprises of 21 convolutional layers, two recurrent layers, two attention layers, and an output layer. Following common practices in deep learning, we optimize these numbers by gradually adding network layers. We track the increase of the decoder performance using 5-fold cross-validation. The performance converges as we stack more residual blocks, LSTM layers, and attention blocks. Additional layers beyond this point tend to cause over-fitting.

We train 15 AI models separately where each model decodes the trajectory of an individual DOF. All 15 models have the same architecture (i.e. number of layers, parameters, etc.) and take the input from all 16 recording channels. There are two technical reasons for this. First, the training process (i.e. learning rate, decay rate, number of epochs, etc.) can be optimized for individual DOF to achieve the best prediction accuracy while preventing over-fitting. For example, here the D5 model (thumb) tends to converge in fewer epochs than the D1 model (little finger) because the signals are more distinguishable on the median nerve. Second, the system would be more robust because if one model has poor accuracy or fails during normal operation, it would not affect the performance of others.

#### Convolutional layers

Convolutional layers are used to encode local patterns of the input. All convolutional layers have 32 filters with a kernel width of three. After each convolutional layer, we apply batch normalization and a rectified linear unit (ReLU) activation. The batch normalization is a technique that normalizes the inputs of a layer to result in a mean output activation of zero and standard deviation of one [25]. It can improve the performance and stability of neural networks. The ReLU activation function which is defined as *f* (*x*) = *max*(0, *x*) is the most popular activation function for deep neural networks [35]. It overcomes the vanishing gradient problem, allowing networks to learn faster and perform better. Inspired by the design of residual networks [21], we stack 10 residual blocks with two convolutional layers each, following a convolutional layer that reduces the input dimensionality. Every other residual block subsamples its inputs by a factor of two. We also apply dropout [50] between residual blocks with a probability of 0.1.

#### Recurrent layers with attention

To accumulate local information across time, we feed the convolutional layer outputs into a two-layer recurrent network, one temporal step at a time. LSTM cells are used as the basic RNN unit. Each LSTM cell has 64 hidden units. After each recurrent layer, we apply a dropout with a probability of 0.2. The recurrent layer outputs are linearly combined across time, with dynamic combination weights determined with a self-attention mechanism. The self-attention block is composed of two dense layers with a ReLU in between, which computes one attention weight for each temporal step. The softmax-normalized weights are multiplied with the hidden states of the LSTM, generating a 64-dimensional code that represents the entire input signal.

#### Output layer

Since our experiments only consist of simple finger motion, a linear projection output is sufficient to decode the hidden code into an output motion trajectory. For more complex motion patterns, this linear decoder can be replaced with a deeper network structure.

### 2.6. AI Models Training and Benchmarking

#### Training process

The dataset is randomly split across all the trials with 50% of the data used for training and 50% used for validation. The validation data are not seen by the model during the training process and do not overlap with the training data. We train the model using a loss function that combines both mean squared error (MSE) and variance accounted for (VAF), which is denoted as:

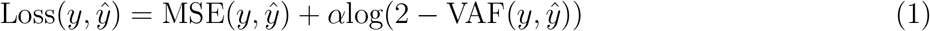

where *α* = 0.05, *y* is the ground-truth trajectory, 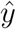 is the estimated trajectory. The values of *y* and 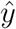 are normalized in a [0, 1] range, where 0 represents the resting position and 1 represents the fully flexing position.

The model parameters are randomly initialized as described by [20]. We use Adam optimizer [31] with the default parameters *β*_1_ = 0.99, *β*_2_ = 0.999, and a weight decay regularization *L*_2_ = 10^−5^. The minibatch size is set to 38 with each training epoch consists of 10 mini-batches. The learning rate is initialized to 0.005 and reduced by a factor of 10 when the training loss stopped improving for two consecutive epochs. These hyperparameters are chosen via a combination of grid search and manual tuning.

#### Baseline models

The proposed deep learning-based AI model is compared with support vector machine (SVM), random forest (RF), multilayer perceptron (MLP), and convolutional neural network (CNN). The SVM parameter is set to *C* = 1. The RF model has 40 trees with a maximum depth of three. The MLP has three hidden layers of 128 units each. The CNN model is created by replacing the recurrent and attention layers of the proposed RNN model with two dense layers of 128 units.

#### Hardware specifications

Training the AI models is a computationally intensive task, thus is done on a desktop PC (Intel Core i7-8086K CPU, NVIDIA Titan XP GPU). On the other hand, data acquisition and motor decoding (i.e. DL inference) require much lower computational resources. We collect training data and perform decoding using both the desktop PC and a laptop (Intel Core i7-6700HQ CPU, NVIDIA GTX 970M GPU) with no significant differences in performance.

#### Performance metrics

We measure the quantitative regression prediction of the decoding algorithm using two metrics MSE and VAF that have the following formulae:

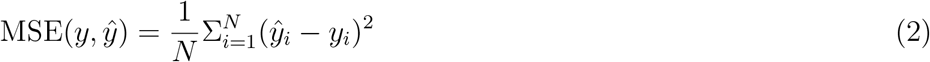

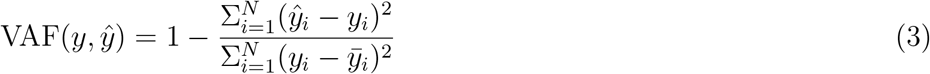

where *N* is the number of samples, where *y* is the ground-truth trajectory, 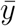 is the average of *y*, and 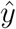 is the estimated trajectory.

## 3. Results

### 3.1. Rich Dynamics of Compound and Single-axon Nerve Activities in ENG Recordings

We use two Scorpius devices to acquire ENG signals from 16 differential electrodes across four implanted arrays. The top four electrodes with the best signal-to-noise ratio (SNR) are selected from each array. We want to include signals from a wide selection of nerve fascicles even though the “sensory” channels may have fewer activities than “motor” channels. A customized adapter is used to manually route recording channels to desirable electrodes. Fig. 6 presents a typical data sequence recorded during an experiment session. Here the patient is asked to perform a mirrored bilateral task of opening and closing all five fingers (M9: fist/grip) on both able and injured (phantom) hands. ENG signals are acquired by the Scorpius system from the injured hand while the intended movements are simultaneously captured by the data glove from the able hand. Fig. 6(A, B) show the filtered data in the low (30-600 Hz) and high (300-3000 Hz) frequency band corresponding to the trajectory of a finger obtained by the data glove. 8-order Butterworth filters are used with the forward-backward zero-phase filtering technique to avoid any distortion. Fig. 6(D, E) show the zoom-in of the data window marked in 6(A, B).

**Figure 6.**
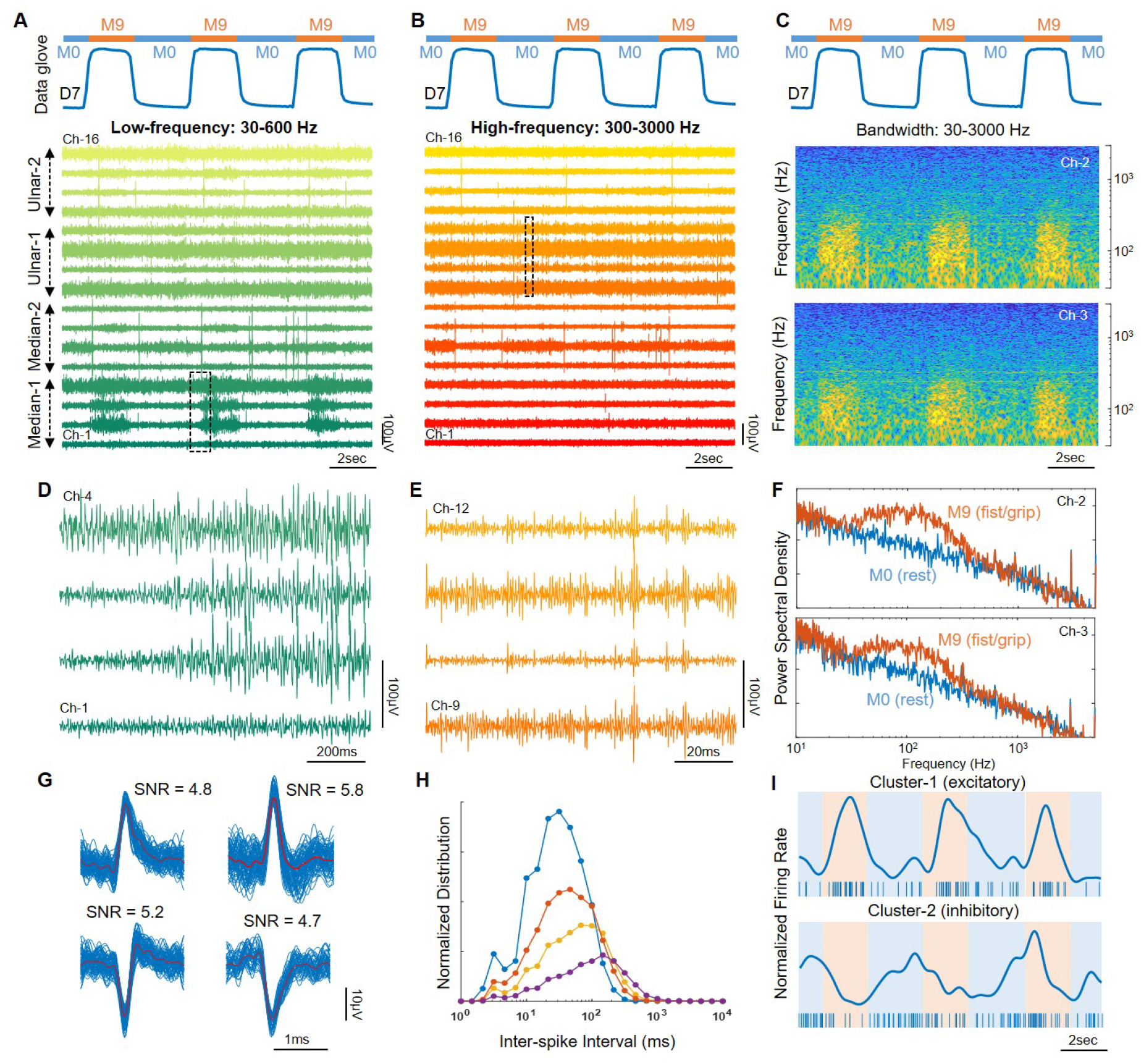
ENG signals acquired by the Scorpius system show rich dynamics in both low- and high-frequency components which are likely associated with nerve compound action potentials (CAP) and single-axon potentials (spikes) respectively. The patient is asked to open (M0) and close (M9) all five fingers repeatedly. (A) Low-frequency (30-600 Hz) and (B) high-frequency (300-3000 Hz) components of the recordings. (C) Spectrogram of two channels what show clear correlation with intended movements. (D, E) Zoom-in of the data window marked in (A, B). (F) Power spectral density analysis of two channels in (C) shows the vibrant differences in ENG dynamics between two gestures M0 and M9. (G) Spike sorting analysis indicates ENG signals in the high-frequency band can be assembled into spike clusters with the characteristics consistent with single-axon potentials. (H) The statistical distribution of the inter-spike interval of different spike clusters. (I) Firing rate of two spike clusters where one cluster is in phase and the other is out of phase with a joint movement.

The acquired ENG signals show rich dynamics of nerve activities which include both compound action potentials (CAP)‡ and single-axon potentials (spikes). CAP is likely associated with the low-frequency band (Fig. 6(A, D)) while spikes tend to be more prominent in the high-frequency band (Fig. 6(B, E)). The proportion of each type of ENG signals on a particular electrode depends on its placement during the implantation surgery and cannot be easily controlled. To maximize the chance of extracting information encoding motor intention, we specifically choose a mixture of electrodes that express both types of data.

Fig. 6(C) presents the spectrogram of two channels (channel 2 and 3) that have particularly large CAP activities. The signal’s dynamics show a clear correlation with the intended movements and distinguishable from the noise floor. Next, we divide the data into segments of resting (M0), movement (M9); and perform the power spectral density estimation using Welch’s method. As shown in Fig. 6(F), there are vibrant differences in ENG dynamics between two gestures M0 and M9.

To evaluate the characteristics of the high-frequency band data, we perform spike sorting analysis using the amplitude-based detection and principal component analysis (PCA) -based sorting techniques as in [67]. All intrafascicular channels display clear neural spikes activities. Fig. 6(G) presents examples of the spike cluster isolated from one recording channel over the entire experiment session (60 min). Fig. 6(H) shows the distribution of inter-spike intervals (ISI). The results show standard waveform and statistics that are consistent with single-axon nerve activities. The spike clusters have the peak-to-peak amplitude ranging from 20-40 *μ*V (SNR = 4-6) which is substantially smaller than CAP signals yet can still be distinctly separated from the noise floor. The noise floor is calculated by taking the rms value of data sequences during resting where no neural activities are present. This results in a 5-6 *μ*Vrms noise floor in 300-3000 Hz which is typical for the electrical noise of the FS amplifier plus the electrode interface. Furthermore, Fig. 6(I) shows the firing rate of two spike clusters modulated by the finger movement. One cluster is in phase and the other is out of phase with a joint movement.

### 3.2. Distinct Spectro-Temporal Signatures Among Different Hand Gestures

Fig. 7 compares the spectrogram of four channels during three different training sessions including thumb bend (M1), ring bend (M4), and fist/grip (M9). Two channels (Ch-2, Ch-3) are from the median nerve while the others (Ch-14, Ch-15) are from the ulnar nerve. The results indicate that each hand gesture has a distinct spectro-temporal signature that is highly correlated with the finger movements. In this study, these signatures are used as the basis to decode the patient’s motor intention. Here we specifically select the channels and gestures with visually observable signatures. In other training sessions and channels, the discrepancy is dimensionally complex and not always obvious to the naked eyes. Therefore, machine learning is the most advantageous approach to implement the motor decoder.

**Figure 7.**
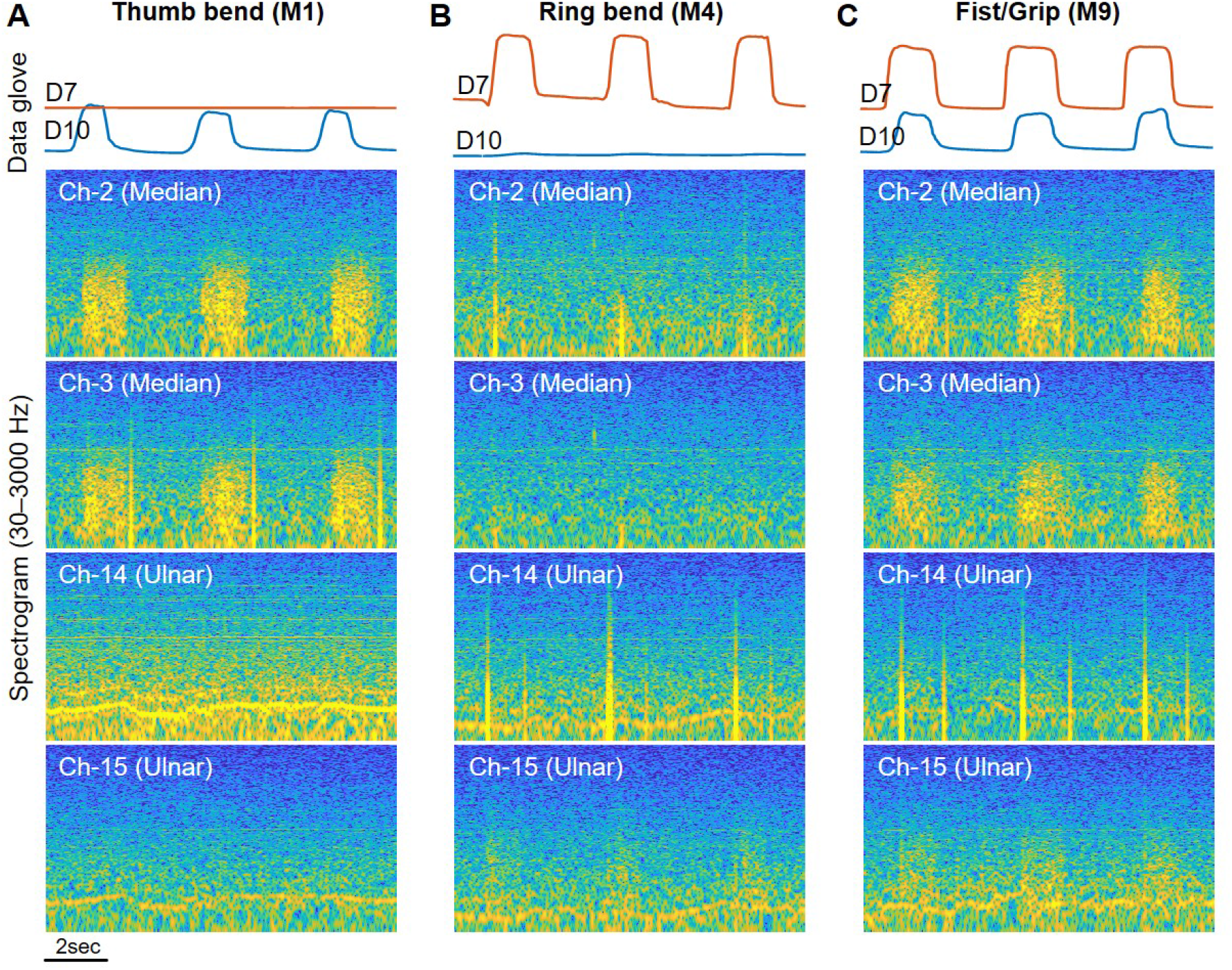
Spectrogram analysis shows unique signatures associated with each hand gesture. Spectrogram of four channels (two from the median nerve, two from the ulnar nerve) during different training sessions is analyzed. Each hand gesture has a distinct spectro-temporal signature that could be visually observed. Nerve activities when bending the thumb (M1 and M9) are more prominent in the median nerve and less notable in the ulnar which is consistent with human anatomy. Opposite observations can be seen with the ring finger (M4 and M9). (A) Spectrogram during thumb bending (M1) training session. (B) Spectrogram during ring finger bending (M4) training session. (C) Spectrogram during fist/grip (M9) training session.

Furthermore, nerve activities are consistent with the human anatomy. The median nerve mostly innervates the thumb, index, middle, and half the ring fingers; while the ulnar nerve is with half the ring and the little finger. As shown in Fig. 7(A, C), when bending the thumb (M1, M9), ENG signals are more prominent in the median nerve and less notable in the ulnar nerve. Opposite observations can be seen with the ring finger (M4, M9) in Fig. 7(B, C). This further reinforces the capability of the Scorpius system in acquiring ENG signals with the necessary information to decode the intended movement of individual fingers.

### 3.3. Dexterous Control of Individual Fingers of A Prosthetic Hand

The performance of the proposed RNN model is compared to other baseline algorithms. They include three “classic” machine learning algorithms: SVM, RF, MLP; and a deep learning model with a simpler architecture: CNN. Fig. 8(A) shows examples of predicted trajectories of 15 DOF produced by each algorithm against the ground-truth movements. In most cases, all algorithms can predict the course motion of each DOF, i.e. whether a finger moves in a specific gesture. However, the two deep learning models, CNN and RNN, always yield a more accurate estimation of the fine trajectory, which is crucial for achieving natural movements.

**Figure 8.**
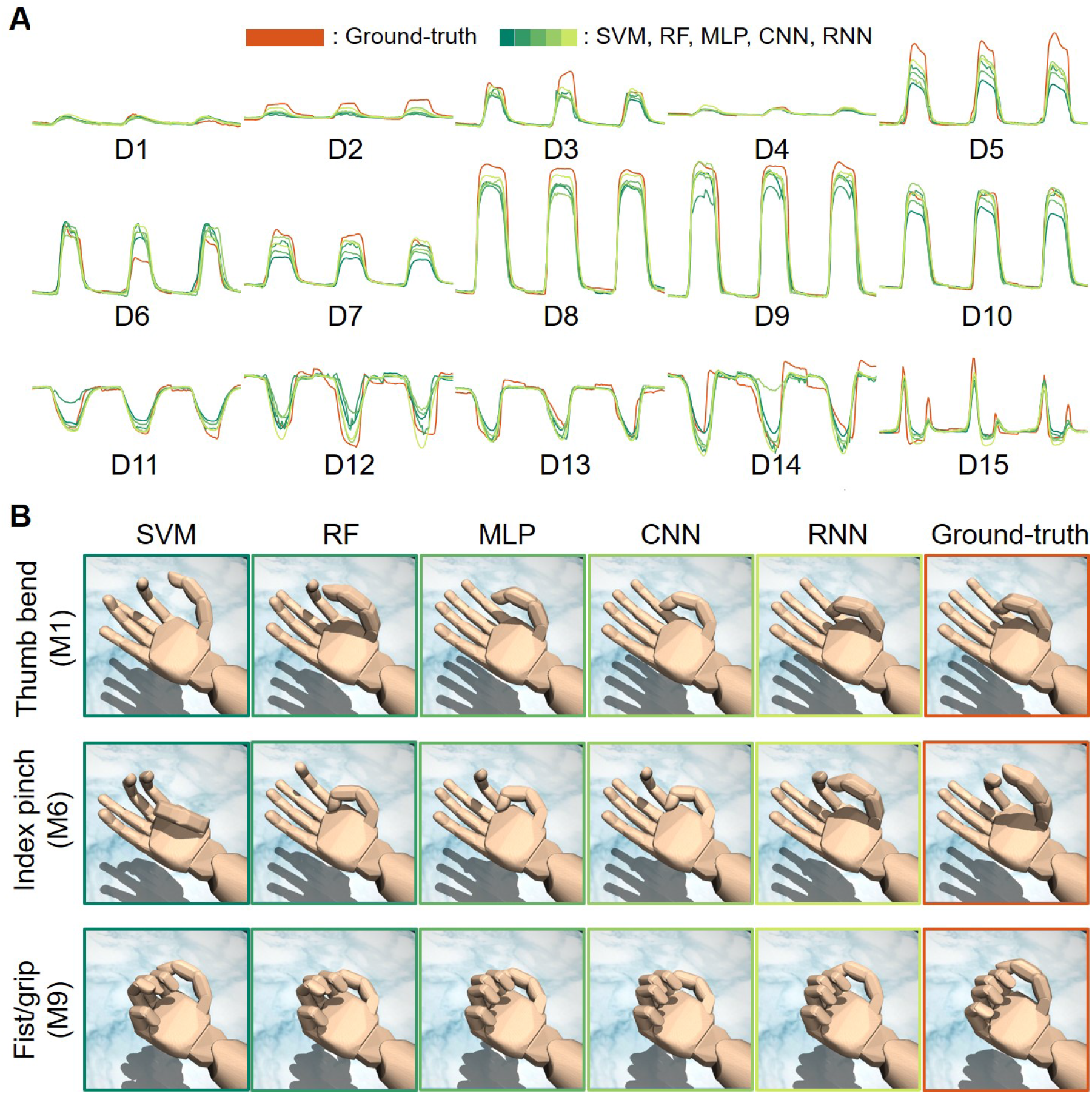
Decoding output of the proposed model in comparison with other machine learning algorithms. We benchmark the proposed RNN model against three “classic” machine learning algorithms: support vector machine (SVM), random forest (RF), multi-layer perceptron (MLP); and a deep learning model with a simpler architecture: the convolutional neural network (CNN). The fine timing and relative relation between multiple DOF are essential to form functional and natural movements. (A) Ground-truth and predicted trajectories of each DOF. (B) The output trajectories are visualized by mapping to a virtual hand (MuJoCo).

In Fig. 8(B), for better visualization, the predicted trajectories are mapped onto a 3D model of the prosthetic hand via the MuJoCo platform [32]. We present the mapping of three gestures: thumb bend (M1), index pinch (M6) and fist/grip (M9) which represent three different classes of motion where one, two and all fingers move respectively. In more complex gestures like M6 and M9, the fine timing and relative relation between multiple DOF become essential to form functional and natural movements. Classic algorithms like SVM, RF, and MLP fail to accomplish this goal even though their raw output trajectories (Fig. 8(A)) may not look much different from the ground-truth.

Fig. 9 reports the comparison of all algorithms using standard quantitative metrics including MSE and VAF. For each DOF, the result shows average performance across all movements. To verify the significance of differences between RNN and the other decoders, we conduct paired t-tests with a Bonferroni correction. The results indicate that the performance differences in MSE and VAF are all statistically significant with *p* < 0.001. The proposed RNN model consistently outperforms other algorithms including CNN across all 15 DOF, yielding measurable lower MSE and higher VAF scores. It is worth noting that the MSE plot is shown on a log scale. In some cases like D7, D9, and D10, RNN offers 3-4 times lower error than SVM and RF. However, MSE cannot be used to compare the relative performance among different DOF. Some joints like D1-D5 move significantly less than the others resulting in lower MSE baseline. This is due to the limitation of the hand gestures used for training and the implementation of the flex sensors in the data glove. VAF scores offer a more realistic benchmark. In most DOF with substantial movements (D5-D15), the RNN achieves VAF scores of 0.7-0.9 (out of a maximum score of 1). These are equivalent to near-natural movements of individual fingers.

**Figure 9.**
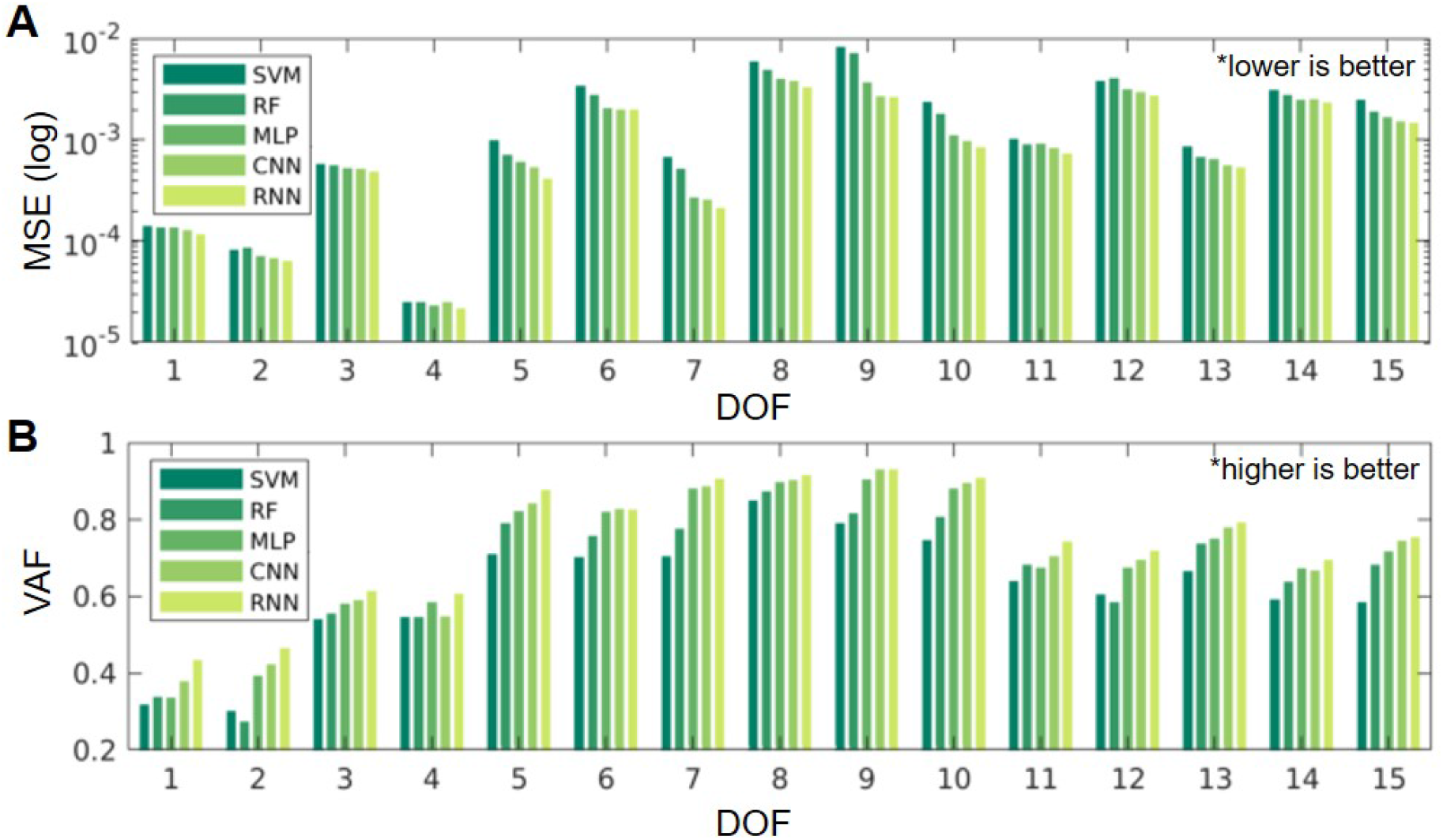
Quantitative metrics showing the proposed model consistently outperforms other algorithms. (A) Mean-square-error (MSE). (B) Variance accounted for (VAF).

## 4. Discussion

Many advanced prosthetic hands available today lack an effective and intuitive control scheme due to the limited accessibility to the human nervous system. Here we present a bioelectric neural interface that could help bridge this gap by tapping into the human mind through the peripheral neural pathways, creating a viable approach towards a direct and intuitive control strategy for multi-DOF prosthetic hands with near-natural dexterity. The platform’s capabilities are clearly demonstrated with empirical evidence through a proof-of-concept motor decoding experiment on a transracial amputee.

### 4.1. Open the Information Conduit Through Peripheral Nerves

The proposed interface opens up a conduit allowing direct access to the nervous system through peripheral nerves. Recorded neural data show characteristics that are consistent with ENG signals originated from bundles of neuron axons in a nerve fascicle. Acquired ENG signals have rich dynamics in both low- and high-frequency bands that are conforming with the compound (CAP) and single-axon (spikes) nerve activities. Those activities are also shown to have distinct spectro-temporal signatures associated with each hand gesture and are consistent with the anatomy/function of the median and ulnar nerve.

The data accessible from this conduit are shown to contain sufficient information contents to enable dexterous control of a prosthetic hand. Motor decoding results show the proposed AI model (RNN) is able to translate the neural patterns into individual finger movements with up to 15 DOF simultaneously. The RNN model outperforms not only classic machine learning algorithms (SVM, RF, MLP) but also the CNN model across all DOF in both performance metrics (MSE, VAF). This implies the motor control signals embedded in the neural data are comprehensive, yet highly complex and intricate. When visualizing through a virtual prosthetic hand, the more fine-detailed prediction of finger trajectories produced by the proposed AI model is shown to be essential to enable lifelike dexterity.

### 4.2. EMG/ENG Nature of Acquired Neural Data

There is an open question that calls for further investigation into the EMG/ENG nature of the acquired neural data. While both EMG and ENG recordings may be used to control basic prosthetic functions such as grasping, the EMG component alone cannot be used to facilitate intuitive hand movements because they simply do not contain any information of missing muscles. Thus a dexterous prosthesis approaching a natural hand is only possible if the decoder has access to the nervous system through the ENG component of the acquired neural data. An ENG-system can also be applied to a wider population of patients with various levels of amputation.

In the current experiment setup, while it is difficult to separate the EMG and ENG components due to their overlapping frequency bandwidth, there is adequate evidence to suggest that the acquired data contain all or mostly ENG signals:

- (i) The recording channels are physically very closed to each other and strategically located far away from residual muscles. The intrafascicular contacts are 0.5mm apart, and the electrode arrays are implanted about 1cm apart. The implantation location is near the far end of the amputated limb which is away from active muscles in the forearm (Fig. 3). As a result, volume-conducted EMG signals should appear on all recording channels with similar waveform and amplitude, which is not evident in the acquired data (Fig. 6 and Fig. 7).
- (ii) ENG components typical of spontaneous single-axon action potentials (i.e. spikes) can be found across all recording channels as shown in Fig. 6, which indicates the electrode contacts successfully interface with nerve fibers. These neural spikes can form clusters with nominal waveform and inter-spike interval. Some spike clusters have excitatory behavior while others have inhibitory behavior.
- (iii) There is a clear separation between control signals in the median and ulnar nerve. This characteristic can only be explained if ENG is the dominant component of the neural data. For example, in Fig. 7, the signals controlling the flexing of the thumb appear strongly in the median channels, but none can be found in the ulnar channels.
- (iv) The AI decoder can predict the movement of all five fingers with high accuracy and dexterity including flexion/extension and abduction/abduction, which suggests the data consist of mostly ENG signals. EMG signals do not contain sufficient information for such tasks because the muscles in the hand are no longer present.
- (v) The patient is asked to purposely contract the residual muscles in the amputated limb but could not reproduce the recorded waveform. We also note that the signals acquired when flexing each finger remain the same regardless of the arm’s position.

### 4.3. Towards Real-time Motor Decoding and Sensory Feedback

The future works will involve developing the existing framework into an online paradigm that is capable of real-time decoding neural data and controlling a physical prosthetic hand. This will not only be a definite demonstration of the proposed platform in practice but also one step closer to a comprehensive solution for clinical uses. The incorporation of real-time control will give the essential visual feedback to the amputees, allowing their brain to adapt the neural pattern to the deep neural network’s behavior, thus further enhancing the decoding accuracy as well as the perception of the hand. The key challenge will be optimizing the components of the platform to work seamlessly together as well as minimize the processing latency.

Furthermore, we are also working on incorporating electrical stimulation capabilities into the proposed platform. The feasibility of using the same electrode array for stimulation has been explored in [40]. This will allow the system to provide neural feedback to the patient including tactile and proprioceptive responses, which are essential for a realistic neuroprosthesis [13, 14]. These features could be facilitated by the Neuronix chip’s support for simultaneous recording and stimulation [65, 67], as well as advances in high-precision, charge-balanced neural stimulators [37, 66].

## 5. Conclusion

We present a technological platform underlying a new class of bioelectric neural interface that is capable of establishing an efficient information conduit between human minds and machines through the peripheral neural pathways. It is achieved thanks to the groundwork in LIFE-FAST microelectrodes, FS-based neural recorders, and deep learning-based AI neural decoder. Empirical data resulted from a pilot human experiment show that the proposed interface could potentially allow amputees to control their prosthetic hands with exceptional dexterity and intuitiveness. Together, these results demonstrate that bioelectric neural interfaces with fully-integrated microelectrodes, bioelectronics, and AI decoders could be the basis for the future human-machine symbiosis.

## Acknowledgments

The surgery and patients related costs were supported in part by the DARPA under Grants HR0011-17-2-0060 and N66001-15-C-4016. The human motor decoding experiments, including the development of the prototype, was supported in part by MnDRIVE Program and Institute for Engineering in Medicine at the University of Minnesota, in part by the NIH under Grant R01-MH111413-01, in part by NSF CAREER Award No. 1845709, in part by Fasikl Inc., and in part by Singapore Ministry of Education funding R-263-000-A47-112, and Young Investigator Award R-263-000-A29-133.

Zhi Yang is co-founder of, and holds equity in, Fasikl Inc., a sponsor of this project. This interest has been reviewed and managed by the University of Minnesota in accordance with its Conflict of Interest policy.

## Footnotes

‡ CAP conventionally refer to the response produced by thousands of individual nerve fibers to electrical stimulation. Because action potentials travel at different velocity on each fiber, the compound potential (CAP) has a broad waveform that contains lower frequency components than individual potential (spike). Here we use the term CAP to refer to a similar modality of neural signals that is a mixture of incoherent individual fiber action potentials originating from the spinal cord and traveling to the muscles at different velocities.

